# Population genome sequencing of the scab fungal species *Venturia inaequalis*, *Venturia pirina*, *Venturia aucupariae* and *Venturia asperata*

**DOI:** 10.1101/477216

**Authors:** Bruno Le Cam, Dan Sargent, Jérôme Gouzy, Joëlle Amselem, Marie-Noëlle Bellanger, Olivier Bouchez, Spencer Brown, Valérie Caffier, Marie De Gracia, Robert Debuchy, Ludovic Duvaux, Thibaut Payen, Mélanie Sannier, Jason Shiller, Jérôme Collemare, Christophe Lemaire

**Affiliations:** IRHS, INRA, Université d’Angers, Agrocampus-Ouest, SFR 4207 QuaSaV, 49071, Beaucouzé, France; Research and Innovation Centre, Fondazione Edmund Mach, via Mach 1, 38010, San Michele All’adige, TN, Italy; UMR 2594, Laboratoire Interactions Plantes Micro-organismes [LIPM], INRA, CNRS, Castanet-Tolosan, France; Unité de Recherche Génomique Informatique [URGI], INRA, Université Paris-Saclay, 78026, Versailles, France.; US1426, INRA, GeT-PlaGe, Genotoul, Castanet-Tolosan, France.; UMR 8298, Institute for Integrative Biology of the Cell (I2BC), CEA, CNRS, Université Paris-Sud, 91405 Orsay, France; UMR 8621, Institut de Génétique et Microbiologie CNRS, Université Paris-Sud, 91405 Orsay, France; Westerdijk Fungal Biodiversity Institute, 3584 CT, Utrecht, The Netherlands

**Keywords:** Venturia, Fusicladium, formae specialis, transposable elements, effectors, apple scab, apple, pear

## Abstract

The *Venturia* genus comprises fungal species that are pathogens on *Rosaceae* host plants, including *V. inaequalis* and *V. asperata* on apple, *V. aucupariae* on sorbus and *V. pirina* on pear. Although the genetic structure of *V. inaequalis* populations has been investigated in detail, genomic features underlying these subdivisions remain poorly understood. Here, we report whole genome sequencing of 87 *Venturia* strains that represent each species and each population within *V. inaequalis*. We present a PacBio genome assembly for the *V. inaequalis* EU-B04 reference isolate. The size of selected genomes was determined by flow cytometry, and varied from 45 to 93 Mb. Genome assemblies of *V. inaequalis* and *V. aucupariae* contain a high content of transposable elements (TEs), most of which belong to the Gypsy or Copia LTR superfamilies and have been inactivated by Repeat-Induced Point mutations. The reference assembly of *V. inaequalis* presents a mosaic structure of GC-equilibrated regions that mainly contain predicted genes and AT-rich regions, mainly composed of TEs. Six pairs of strains were identified as clones. Single-Nucleotide Polymorphism (SNP) analysis between these clones revealed a high number of SNPs that are mostly located in AT-rich regions due to misalignments and allowed determining a false discovery rate. The availability of these genome sequences is expected to stimulate genetics and population genomics research of *Venturia* pathogens. Especially, it will help understanding the evolutionary history of *Venturia* species that are pathogenic on different hosts, a history that has probably been substantially influenced by TEs.

## Introduction

Fungal species from the *Venturia* Sacc. genus (anamorph is *Fusicladium* Bonord.; Beck *et al*., 2005) are pathogenic on important *Rosaceae* host crops including apple and pear (Sivanesan, 1977). They are responsible for scab diseases which dramatically reduce the economic value of fruits by causing black lesions on the skin (Bowen *et al*., 2011). Taxonomically, scab fungi belong to the *Dothideomycetes* class and to the order *Venturiales* within the *Venturiaceae* family (Zhang *et al*., 2011).

Scab caused by *Venturia inaequalis* Cooke (Wint.) is the main fungal disease on apple, control of which may require over 20 chemical treatments each year. *V. inaequalis* has been extensively studied as a model for pathogens on woody hosts, particularly because it is amenable to *in vitro* crossing (Bénaouf and Parisi, 2000). *V. inaequalis* has been described on several genera of Rosaceous including *Malus*, *Pyracantha*, *Eriobotrya* and *Sorbus* (Sivanesan, 1977; Jones and Aldwinckle, 1990). In some cases, host specificity occurs; for instance strains from *Pyracantha* are not able to infect *Malus* and vice-versa, which is referred to as *formae specialis* f. sp *pyracantha* and f.sp *pomi*, respectively (Le Cam *et al*., 2002). Worldwide sampling of *V. inaequalis* strains has identified divergent populations, mostly reflecting their hosts: wild apple (*Malus sieversii*, *M. orientalis*, *M sylvestris*, *M. floribunda*) and domesticated apple (*Malus x domestica*) trees (Gladieux *et al*., 2008; Gladieux *et al*., 2010a; Gladieux *et al*., 2010b; Leroy *et al*., 2013; Ebrahimi *et al*., 2016; Leroy *et al*., 2016). Certain populations have experienced a secondary contact with subsequent gene flow, resulting in the introduction from a wild population into the agricultural compartment of strains that are virulent on resistant cultivars carrying the *Rvi6* resistance gene (Guérin *et al*., 2007;Lemaire *et al*., 2016). On apple, scab can also be caused by another *Venturia* sp., *V. asperata* initially described as a saprophyte (Samuels and Sivanesan, 1975), and recently reported as a pathogen on apple trees (Caffier *et al*., 2012). Other *Venturia* species cause scab damages on their respective *Rosaceae* hosts, *V. pirina* (Aderh.) on European pear (*Pyrus communis*), *V. nashicola* on Asian pear (*Pyrus pyrifolia*), and *V. carpophila* on apricot (Crous *et al*., 2007; Sivanesan, 1977). Other scab fungi are pathogenic on various tree hosts, including *Fusicladium effusum* on pecan, *F. oleaginum* on olive, *V. populina* on poplar, *V. saliciperda* on willow and *V. fraxini* on ash, among many others (Sivanesan, 1977; Zhang *et al*., 2011; Young *et al*., 2018). Despite being economically important pathogens, the genetic basis of the interactions between scab fungi and their respective hosts remains poorly understood (Singh *et al*., 2005; Steiner and Oerke, 2007; Kucheryava *et al*., 2008; Bowen *et al*., 2009; Mesarich *et al*., 2012; Cooke *et al*., 2014; Shiller *et al*., 2015; Deng *et al*., 2017).

Genome sequences are already available for *V. inaequalis f. sp. pomi, V. inaequalis f. sp. eriobotrya*, *V. pirina* and *F. effusum*, and they revealed that *V. inaequalis* strains tend to have a higher repeat content compared to other species, with up to 47% of their genome sequence covered by repetitive elements (Shiller *et al*. 2015; Bock *et al*., 2016; Deng *et al*., 2017; Passey *et al*., 2018). These studies highlighted in *Venturia* genomes the expansion of putative effector gene families, genes that are known to determine plant-fungus interactions (Shiller *et al*., 2015; Deng *et al*., 2017). The high repeat content and the employed Illumina sequencing strategy resulted in assemblies with between 1,012 and 1,680 scaffolds (Shiller *et al*. 2015; Deng *et al*., 2017) and up to 231 contigs using the PacBio technology (Passey *et al*., 2018). Genetic studies need higher quality genome assemblies and genomic population studies require re-sequencing of many strains from diverse populations.

Here, we report the PacBio genome assembly of the *V. inaequalis* EU-B04 reference strain and re-sequencing of 79 strains from two *formae specialis* and from diverse populations within *V. inaequalis f. sp. pomi*. We also present re-sequencing of *V. pirina* strains and whole genome sequencing of *V. aucupariae* and *V. asperata*. Availability of these genome sequences will greatly facilitate genetic and genomic population studies to better understand the evolution of *Venturia* pathogens and their interactions with different hosts.

## Materials and Methods

### Origin of *Venturia* strains

All 87 *Venturia* strains used in this study are described in Table S1. Forty strains of *V. inaequalis* sampled in Kazakhstan on wild apple *M. sieversii* were used, of which half belong to the wild CAM (Central Asian Mountains) population and the other half to the domestic CAP (Central Asian Plains) population as described in Gladieux *et al*., 2010b. Other *V. inaequalis* strains were isolated from *M. x domestica* cultivars that do not carry the *Rvi6* resistance gene (6 domEU and 2 domASIA), from *M. x domestica* cultivars carrying the *Rvi6* resistance gene (5 dom*Rvi6*), from ornamental *M. floribunda* (4 FLO), from wild *M. sylvestris* (6 SYL) and *M. orientalis* (6 ORI), from *Pyracantha* (5 PYR), from *E. japonica* (5 LOQ) and from *Sorbus aucuparia* (1). In addition, strains of *V. pirina* (3), *V. aucupariae* (1) and *V. asperata* (3) were isolated from *Pyrus communis* (pear), *S. aucuparia* and *M. x domestica*, respectively (Table S1).

All strains were grown at 18°C on cellophane membranes displayed on PDA-YE (Potato-Dextrose Agar (BD Difco) supplemented with 0.3% Yeast Extract (Sigma-Aldrich)) plates with a 12h/12h light/night regime. They are stored at -20°C on dried cellophane membranes in the *Venturia* collection of the Institut de Recherche en Horticulture et Semences (IRHS, Beaucouzé, France).

### Genomic DNA extraction, library construction and DNA sequencing

High molecular weight genomic DNA from *V. inaequalis* EU-B04 strain was extracted from mycelium grown on PDA-YE covered with cellophane membrane using the method described in Cheeseman *et al.* (2014) (extraction with CTAB followed by purification on an isopycnic gradient), except that ethidium bromide was used instead of DAPI for the purification of genomic DNA on a cesium chloride gradient. Ethidum bromide was preferred because it resulted in a single band of genomic DNA, whilst DAPI resulted in the dispersion of genomic DNA in several bands of different AT contents.

Genomic DNA quantity and quality were validated at the GeT-PlaGe core facility (Toulouse, France) using Qubit (Thermofisher), Nanodrop (Thermofischer), and Fragment Analyzer (Advanced Analytical) before purification on 0.5X AMPure PB beads and shearing at 40 kb using the Megaruptor system (Diagenode). Libraries were prepared according to the “20-kb Template Preparation Using BluePippin^TM^ (Sage Science) Size-Selection System (15-kb size cutoff)” PACBIO protocol using 10 µg of sheared genomic DNA and the SMRTBell template Prep Kit. DNA fragments from 10 kb to 90 kb were selected. The sequencing run was performed on a PacBio RSII sequencer with the OneCellPerWell Protocol (OCPW) on C4 P6 chemistry and 360 minute movies. Another PacBio sequencing was performed at the Cold Spring Harbour core genomics facility (CSHL, NY, USA); genomic DNA was sheared and purified, and a long-insert library was prepared following the protocol described in Quail *et al*. (2012). The library was sequenced on four SMRT cells using the C3 P6 chemistry using 2x 60 minute movies.

Genomic DNA of all other *Venturia* strains was extracted from mycelium grown on PDA-YE plates using Möller’s protocol (Möller *et al*., 1992). Libraries for CAM and CAP strains were prepared using SPRIworks-Fragment Library System I (Beckman Coulter) and sequenced at GATC Biotech (Konstanz, Germany) on HiSeq 2000 using TruSeq Multiplex Sequencing Primer and TruSeq PE (2×50bp) (Illumina, San Diego, CA). Libraries for the remaining strains were prepared using the TruSeq Nano kit (Illumina, San Diego, CA) and sequenced at the GeT-PlaGe core facility (Toulouse, France) using SBS v3 (2×100 pb) or v4 (2×125 pb) reagent kits and using HiSeq 2000 or HiSeq 2500. For library construction, genomic DNAs were fragmented by sonication and sequencing adaptors were ligated. Libraries were amplified with eight PCR cycles; their quality was assessed using a Bioanalyzer (Agilent) and quantity measured by quantitative PCR using the Kapa Library Quantification Kit (KapaBiosystems).

### Genome assembly

For the PacBio genome, the PBcR wgs8.3rc2 assembly pipeline (Berlin *et al*. 2015) was used to perform read correction and assemble the corrected reads. The high sequencing depth and quality of raw data allowed modifying several parameters to increase stringency: i) only raw reads longer than 5 kb were considered; ii) 50x of corrected reads were collected (assembleCoverage=50); iii) the runCA part of the pipeline was run with a minimum overlap length of 1 kb (ovlMinLen=1000); and iv) the unitigger bogart was tuned to consider overlaps with a very low error rate only (utgErrorRate = 0.0075 and utgMergeErrorRate = 0.04). Spurious contigs that were fully included in larger contigs were removed using a script based on ncbi-megablast to perform the all-*vs*-all analysis, combined with additional filters (minimum identity of 93%, minimum overlap of 500 nt and maximum overhang length of 500 nt). Bacterial contaminant contigs were identified using ncbi-blastx search against the NR database (downloaded in May 2016) and manual inspection. A total of 46 contigs were removed (maximum length of 113 kb) based on best hits with bacterial proteins confirmed with a GC percent analysis. Finally, the genome was polished with quiver (Chin *et al*. 2013) using stringent alignment cutoffs for pbalign (--minLength 3000 --maxHits 1), ensuring a mapping as specific as possible. Assemblies of other genomes were performed by two algorithms: SOAPdeNovo2 (Zerbino *et al*., 2008) and Velvet (Luo *et al*., 2012) adapted for de novo genome assembly using next-generation sequencing (NGS) short reads.

Genome statistics were determined using Quast 2.3 (Gurevich *et al*., 2013). The GC content and distribution in each genome was analysed using OcculterCut (Testa *et al*., 2016). Completeness of genome assemblies was assessed using BUSCO v1.1b1 (Simão *et al*. 2015) and the fungal BUSCO profile (busco.ezlab.org/v1/files/fungi_buscos.tar.gz). The program was run in “genome assembly” mode using the metaparameters of *Fusarium graminearum* for AUGUSTUS gene prediction.

### SNP discovery, false discovery analysis and population structure of *Venturia spp*

For all individuals, genome sequencing reads were cleaned using Trimmomatic 0.36 (Bolger *et al*. 2014) before mapping on the EU-B04 reference genome using BWA-MEM (Li and Durbin 2009). For species other than *V. inaequalis*, read mapping was refined with Stampy 1.0.29 (Lunter and Goodson 2011) using the parameter “-- bamkeepgoodreads” and the parameter “--substitutionrate” was set to 0.15, 0.075, and 0.07 for *V. asperata*, *V. pirina* and *V. aucupariae*, respectively. SNP discovery was performed using the Unified Genotyper module of GATK 3.6 (McKenna *et al*., 2010), applying indel realignment and duplicate removal according to GATK Best Practices recommendations (DePristo *et al*. 2011; Van der Auwera *et al*. 2013). Instead of applying any posterior quality score recalibration and s tandard hard filtering, we performed a SNP false discovery rate (FDR) analysis making use of clones present in our dataset (Table S1). The FDR was estimated by simply counting the number of SNPs for each pair of clones. All steps but the Trimmomatic, Stampy and FDR analysis were integrated in a pipeline set using the TOGGLE toolbox (Monat *et al*. 2015). In order to assess the discrimination power of our resulting true positive SNP dataset, we computed a PCA of all the non-clonal *Venturia inaequalis* strains using PLINK v1.90b3.44 (Purcell *et al*. 2007).

### RNA extraction and sequencing

*V. inaequalis* EU-B04, *V. pirina* VP36 and *V. asperata* 2349 strains were grown on seven different agar media: PDA-YE, apple medium (10% organic apple sauce in water), and Vogel media (Vogel, 1956) without carbon or nitrogen source, or supplemented with 1% glucose or 1% saccharose (with and without 80 µM zebularin). Total RNAs were extracted using the kit Nucleospin RNA Plant (Macherey-Nagel, Düren, Germany), according to the manufacturer’s recommendations. Total RNAs from complex media and from defined media were combined in two respective pools. Paired-end 2×150 bp libraries were prepared using the Illumina TruSeq Stranded mRNA kit (Illumina, San Diego, CA) and sequenced at the GeT-PlaGe core facility (Toulouse, France) using a paired-end read length of 2×150 pb with the Illumina HiSeq3000 sequencing kits and using HiSeq 3000. Libraries were prepared according to Illumina’s protocols on a Tecan EVO200 liquid handler. Messenger RNAs were selected using poly-T beads, fragmented to generate double stranded cDNA; and then sequencing adaptors were ligated. Libraries were amplified with ten PCR cycles. Their quality was assessed using a Fragment Analyzer (Advanced Analytical) and quantity measured by quantitative PCR using the Kapa Library Quantification Kit (KapaBiosystems).

### Flow cytometry

Analyses were performed according to Marie and Brown (1993). About 1 cm^2^ of mycelium grown on PDA-YE plate was scraped and resuspended in 500 µL of cold nuclear buffer (45 mM MgCl_2_, 30 mM sodium citrate, 20 mM MOPS pH 7, 0.02% Triton X-100, 1% (w/v) cold polyvinyl pyrrolidone, 5 mM sodium metabisulfite, 0.05 mg/mL RNase (Roche, Basel, Switzerland)). A ligule of *Arabidopsis thaliana* cv. Columbia-0 was added as an internal standard. Samples were cut with a razor blade, filtered with 30 µm CellTrics paper (Sysmex Partec, Kobe, Japan) and stained with 50 µg/mL propidium iodide (Fluka). After 5 min incubation at 4°C, cytometric analysis was performed on 20,000 nuclei using a CyFlo3 cytometer (Sysmex Partec, Kobe, Japan). Intensity ratio between 1C fungal nuclei and 2C (= 0.33 pg, corresponding to 326 Mbp) *A. thaliana* nuclei was determined to measure the size of fungal genomes (Dolezel *et al*., 2003).

### Gene and transposable element prediction and annotation

Genes of the EU-B04 reference genome were predicted using the Braker1 pipeline (Hoff *et al*., 2016). RNA-alignments to the EU-B04 genome from all the *in vitro* conditions described above were used to provide extrinsic hints for gene prediction in this pipeline. Genes in all other genomes were predicted using AUGUSTUS version 3.2.3 (Stanke *et al.,* 2008) utilizing the species training parameter files created in the Braker1 pipeline above.

Secreted proteins in the predicted proteome were annotated with SignalP version 4.1 (Petersen *et al*., 2011) using default parameters. Small-secreted proteins (SSPs) were identified as secreted proteins with a maximum of 300 amino acids, containing no Glycosylphosphatidylinositol (GPI) anchor motif according to Big-PI (default parameters; Eisenhaber *et al*., 2004) and PredGPI (False-Positive rate lower than 1e-3; Pierleoni *et al*., 2008), and no transmembrane domain in the mature form according to TMHMM (default parameters; Krogh *et al.*, 2001).

The REPET pipeline (urgi.versailles.inra.fr/Tools/REPET) was used for the detection, classification (TEdenovo; Flutre *et al*., 2011; Hoede *et al*., 2014) and annotation (TEannot; Quesneville *et al*., 2005) of transposable elements (TEs). The TEdenovo pipeline was used to search for repeats in *V. inaequalis* EU-B04 genome sequence. The first step used Blaster with the following parameters: identity > 90 %, HSP (high-scoring segment pairs) length >100 bp and <20 kb, e-value ≤ 1e-300. HSPs were clustered using three different methods: Piler (Edgar and Myers, 2005), Grouper (Quesneville *et al*., 2003) and Recon (Bao and Eddy, 2002). The MAP software (Huang, 1994) was used to make multiple alignments of the 20 longest members of each cluster containing at least 3 members and build a consensus. Consensus sequences were then classified on the basis of their structure and similarities relative to Repbase Update (v17.11) (Jurka *et al*., 2005) and PFAM domain library v2 (Finn *et al*., 2014), before redundancy removal with Blaster and Matcher as performed in the TEdenovo pipeline. Consensus sequences with no known structure or similarity were classified as “unknown”.

### Phylogenetic species tree

All strains of *V. pirina*, *V. asperata* and *V. aucupariae*, and one isolate from each population of *V. inaequalis* were used to build a phylogenetic tree. The *Dothideomycetes Zymoseptoria tritici* IPO323 and *Alternaria alternata* SRC1lrK2f (genome.jgi.doe.gov) were used as outgroups. The DNA sequence of β-tubulin *TUB1* (KOG1375), eukaryotic initiation factor 3, subunit b *eIF3b* (KOG2314), phosphoadenylphosphosulphate reductase *PAP* (KOG0189) and ribosomal protein S9 *RPS9* (KOG1753) genes were retrieved in all genomes and each were aligned using MUSCLE (Edgar, 2004). Regions with low quality alignment were removed using Gblocks (default parameters; Castresana, 2000) prior to concatenation of the four alignments. A maximum-likelihood tree was built with MEGA6 (Tamura *et al*., 2011) using the best-fit Tamura-Nei substitution model with 5 gamma distribution categories and 500 bootstrap replicates.

### Data availability

All genome sequencing data generated in this study are accessible (assembly fasta files) in GenBank under the BioProject number PRJNA407103.

## Results and Discussion

### Phylogeny of *Venturia* species

Phylogenetic relationships between the different *Venturia* species were determined using the *Dothideomycete* species *Zymoseptoria tritici* and *Alternaria alternata* as outgroup to the *Venturiales* order (Zhang *et al*., 2011; Bock *et al*., 2016). The maximum likelihood tree shows that *V. asperata* is basal to the other *Venturia* species (Figure 1A). *V. aucupariae* has diverged more recently from *V. inaequalis* than *V. pirina*. Phylogenetic relationships among *V. inaequalis* strains from different populations cannot be accurately resolved, probably because of the existence of incomplete lineage sorting and gene flow between populations (Gladieux *et al*., 2010b; Lemaire *et al*., 2016; Leroy *et al*. 2016).

**Figure 1.**
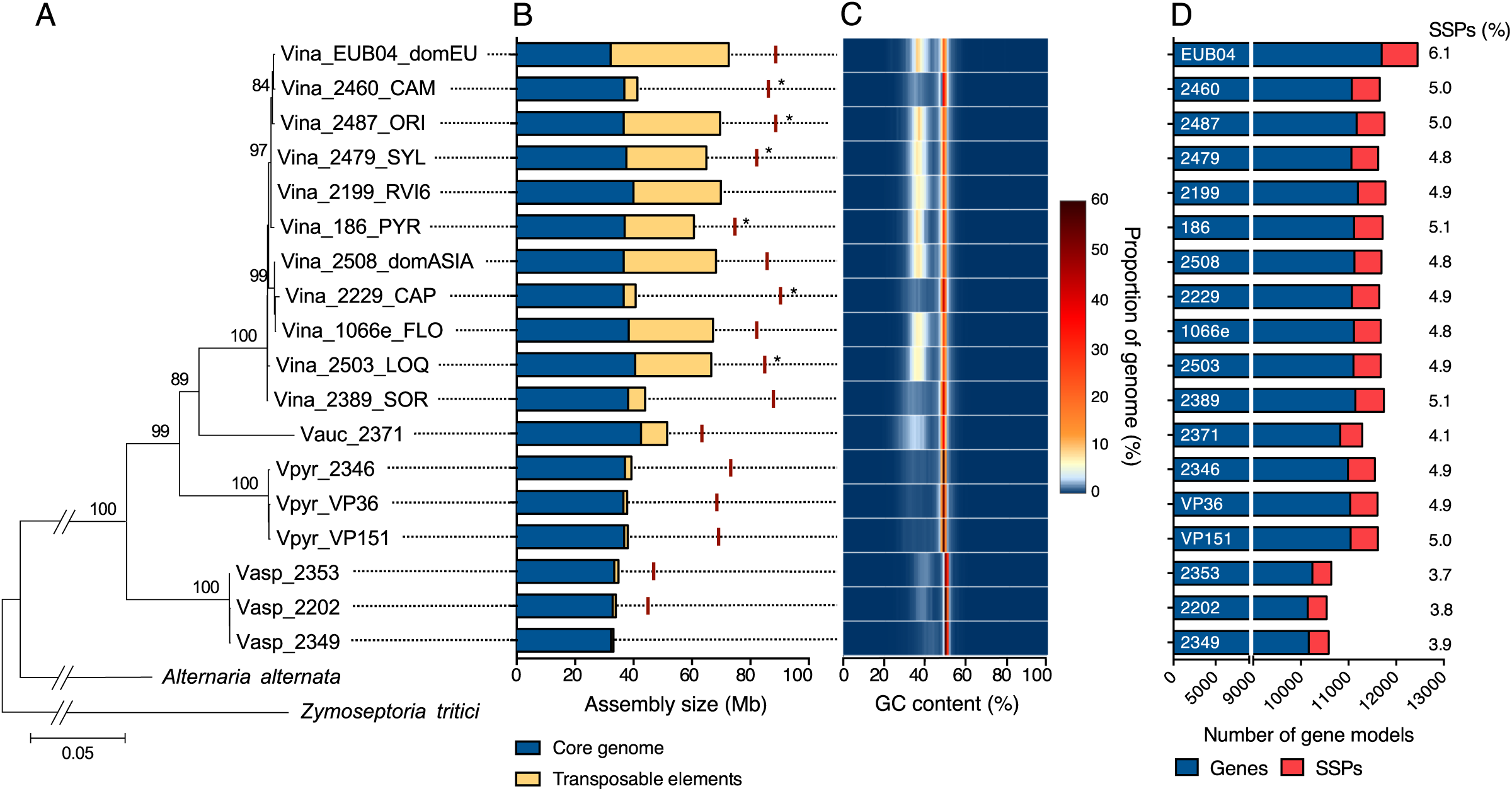
Phylogeny of *Venturia* species and genome characteristics. A) A four-gene maximum-likelihood phylogenetic tree was built using the *Dothideomycete* species *Alternaria alternata* and *Zymoseptoria tritici* as outgroup. Numbers on the branches indicate bootstrap values. The name of the isolates indicates the species (Vina: *V. inaequalis*; Vauc: *V. aucupariae*; Vpir: *V. pirina*; Vasp: *V. asperata*), name in the collection and, for *V. inaequalis* isolates, the population of origin (see Table S1). B) Assembly size is shown for isolates used in the phylogenetic analysis. The size of the assembly covered by transposable elements was determined using the REPET pipeline (urgi.versailles.inra.fr/Tools/REPET). Red lines indicate actual genome sizes as determined by flow cytometry. Stars indicate that the genome size value is the average of other isolates from a given population. C) The GC composition in each genome was determined using OcculterCut (Testa *et al*., 2016). D) Gene models were predicted using Braker1 (Hoff *et al*., 2016). Small-secreted proteins (SSPs) contain a predicted leader signal peptide, do not contain a transmembrane domain or GPI-anchor signal, and are shorter than 300 amino acids. Numbers on the right indicate the fraction of the predicted proteome that is represented by SSPs.

### Genome assemblies

The assembly size of the different *V. inaequalis* strains varies greatly from 30 to 73 Mb, with a median of 7,435 scaffolds (from 66 to 31,301), an N50 median of 55.7 kb and an estimated sequencing coverage ranging from 8.8x to 150x (Figure 1B and Table S1), consistent with previous sequencing of *V. inaequalis* strains (Shiller *et al*., 2015; Deng *et al*., 2017; Passey *et al*., 2018). As expected, the best assembly result was obtained for the EU-B04 reference strain with PacBio sequencing (66 scaffolds, N50 = 2,978 kb). Genomic DNA library construction likely explains the variability in assembly size as certain protocols might have excluded long repetitive regions. It is especially the case of CAM and CAP strains that both exhibit a genome size half the size of genomes of strains found on apple trees. In this case, library construction kits might have filtered the major part of AT-rich regions. The actual genome size of *V. inaequalis* strains is over 80 Mb as measured using flow cytometry (average size of 85.1 ± 5.0 Mb; Figure 1B and Table S1). The measured genome sizes suggest that scaffolds in the PacBio assembly are separated by extremely long stretches of repetitive elements. The average assembly size for *V. aucupariae*, *V. pirina* and *V. asperata* strains is 52, 38 and 34 Mb, respectively (Figure 1B and Table S1). The genome of these species appeared to really contain fewer repeats as library construction and sequencing strategy for these genomes was similar to those of *V. inaequalis* genomes for which repetitive elements were sequenced. However, flow cytometry determined genome sizes of 63.6 ± 1.6 Mb, 70.6 ± 2.7 Mb and 46.1 ± 1.2 Mb for *V. aucupariae*, *V. pirina* and *V. asperata*, respectively (Figure 1B and Table S1). The former two species appear to contain a significant amount of repetitive elements that could not be deduced from their genome assemblies (Shiller *et al*., 2015; Deng *et al*., 2017).

The genome structure of *V. inaequalis* is also a mosaic of GC-equilibrated and AT-rich regions as shown for the EU-B04 reference isolate (Figure 2). Indeed, the GC-content distribution in all genomes exhibits a bimodal pattern typical of genomes with large proportions of AT-rich regions (Figure 1C; Testa *et al*., 2016). These regions cover on average 45.3% of *V. inaequalis* genome assemblies with a size over 60 Mb, while they cover on average 32.5%, 6.6% and 9.9% of *V. aucupariae*, *V. pirina* and *V. asperata* assembled genomes, respectively (Table S1). Considering the measured genome sizes, these values are likely underestimated and AT-rich regions may represent from 30% in *V. asperata* to 50% in *V. pirina*. The GC content of AT-rich regions in *Venturia* species ranges from 33.5% to 41.3% (Table S1), which is similar to the 33.8% reported for *Leptosphaeria maculans* (Testa *et al*., 2016), another species with a mosaic genome structure of GC-equilibrated and AT-rich regions (Rouxel *et al*., 2011).

**Figure 2.**
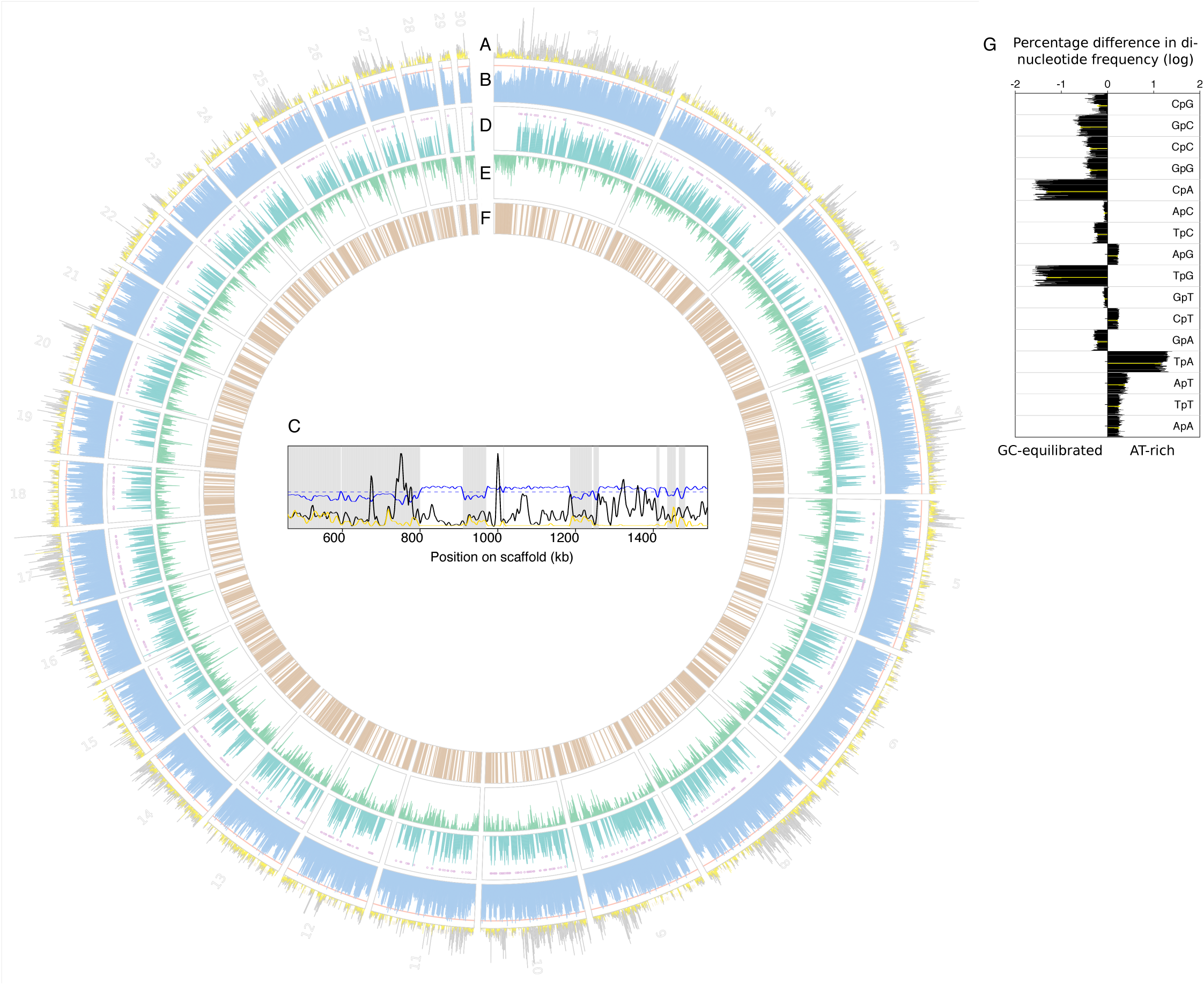
Genome features of the *Venturia inaequalis* EU-B04 reference strain. Twenty-nine out of 66 scaffolds are shown, representing 95% of the total assembly. Scaffold number is indicated on the outer circle. Tracks show (A) Single-Nucleotide Polymorphism (SNP) density between isolates (black) and clones (yellow); (B) GC content (%), the orange line indicate 50%; (C) Gene density according to Braker1 gene prediction, magenta circles indicate the localization of small secreted proteins (SSPs). (D) transposable element density according to REPET analysis. (E) RIP (Repeat Induced Point mutation) index). (F) close-up on scaffold 1 showing SNP densities and GC content as described in (A) and (B) tracks, respectively, the dotted blue line indicating the GC content threshold used by OcculterCut (Testa *et al*., 2016) to separate GC-quilibrated and AT-rich regions; the latter are highlighted with grey shading. (G) Difference in di-nucleotide frequency between GC-equilibrated and AT-rich regions in genomes of all isolates as determined by OcculterCut. Negative and positive values indicate over-representation in GC-equilibrated and AT-rich regions, respectively. Values for the EU-B04 strain reference assembly are indicated in yellow.

Despite the difference in assembly size, the BUSCO assessment of gene content indicates a similar and very high level of completeness for most strains (median of 98% complete single copy genes). This result also indicates that most of the predicted genes are located in GC-equilibrated regions, which represent the most well sequenced and assembled in our dataset. Six strains show a retrieval rate of BUSCO genes lower than 90% (Table S1), meaning that genome sequencing and/or assembly quality is very low and these genomes should be used with caution in further analyses.

### SNP discovery and population structure of *Venturia inaequalis*

Evaluation of the divergence between all strains based on BUSCO genes identified six pairs of strains and one triplet with very short genetic pairwise distances, suggesting they are clones (2225/2446, 2226/2450, 2447/2458, 2474/2475, 2478/2480, 186/2269 and 2503/2504/2263). Consistently, fewer SNPs were discovered between these pairs compared to other strains from the same populations (Table 1). We could use this information to perform a SNP false discovery analysis because we expect *de novo* mutations to be rare between clones of *V. inaequalis* as it reproduces sexually each year (Bowen *et al*., 2011). Our data suggests that 25 to 50% of SNPs identified without masking genomes are false positives (Table 1). About 90% of SNPs identified between clones are located throughout the genome within AT-rich regions that are very rich in transposable elements (Table 1, Figure 2A, 2D and 2F). This observation is most likely explained by the mapping at the same locus of highly similar reads from different genomic origins. In contrast, GC-equilibrated regions appear to contain a higher proportion of SNPs discovered between non-clonal strains, which are enriched on certain scaffolds (Table 1, Figure 2A and 2F). Masking AT-rich regions as determined by OcculterCut eliminated between 88.7% and 96.6% of false positive SNPs for a saving of 16.5% to 48.4% of the initial SNPs between non-clonal strains (Table 1). The proportion of retained SNPs reached 52% in the global dataset of 12.7 million SNPs identified between all strains. The fact that GC-equilibrated regions still contain 3.4% to 11.3% of false positive SNPs is likely explained by the expansion of gene families in the genome of *Venturia* species (Table 1; Shiller *et al*., 2015; Deng *et al*., 2017). This observation suggests that masking AT-rich regions or transposable elements alone often may be not enough to reach an acceptable FDR. As reported here, sequencing two clones or twice the same individuals easily allows detecting regions with high level of FDRs and is of great help to define hard filtering and/or SNP recalibration settings, especially for highly repetitive or polyploid genomes of sexual organisms.

**Table 1.** **Single Nucleotide Polymorphism (SNP) false discovery analysis in *Venturia inaequalis* populations** Isolates originate from populations isolated on loquat (LOQ), firethorn (PYR), *Malus sylvestris* (SYL) and on *Malus sieversii* in Central Asian Mountains (CAM) and Central Asian Plains (CAP).

|  | LOQ population |  |  | CAM population |  |  | CAP population |  |  | SYL population |  |  | PYR population |  |  |
| --- | --- | --- | --- | --- | --- | --- | --- | --- | --- | --- | --- | --- | --- | --- | --- |
|  | 2503 | 2503 | FDR (a) | 2225 | 2225 | FDR (a) | 2475 | 2229 | FDR (a) | 2478 | 2478 | FDR (a) | 186 | 2266 | FDR (a) |
|  | 2504 | 2505 |  | 2446 | 2447 |  | 2474 | 2474 |  | 2480 | 2479 |  | 2269 | 2269 |  |
|  | (c) |  |  | (c) |  |  | (c) |  |  | (c) |  |  | (c) |  |  |
| **Total number of SNPs** | 130,507 | 333,955 | 39.1% | 94,845 | 341,581 | 27.8% | 125,654 | 522,134 | 24.1% | 120,622 | 393,845 | 30.6% | 125,196 | 257,874 | 48.5% |
| **SNPs in AT-rich region (b)** | 120,670 | 201,303 | 59.9% | 88,345 | 205,100 | 43.1% | 116,931 | 269,189 | 43.4% | 113,284 | 328,956 | 34.4% | 118,700 | 196,239 | 60.5% |
| **SNPs in GC-equilibrated regions (b)** | 9,837 | 132,652 | 7.4% | 6,500 | 136,481 | 4.8% | 8,723 | 252,945 | 3.4% | 7,338 | 64,889 | 11.3% | 6,496 | 61,635 | 10.5% |

(a) FDR: False Discovery Rate
(b) GC-equilibrated and AT-rich regions (51.4% and 48.6% of the EU-B04 reference genome, respectively) were determined using OcculterCut (Testa *et al.*, 2016).
(c) indicate clonal isolates

Once cleared of the false positive SNPs, our dataset allowed a clear discrimination of the *Venturia inaequalis* strains according to their host of origin as shown by the PCA (Figure 3). This result is highly congruent with our phylogeny (figure 1A) and previous studies (Gladieux *et al*., 2010,a). It is worth noting though that the individual 2389, sampled on *Sorbus aucuparia* and initially thought as a *V. aucupariae* individual, strongly clusters with the *V. inaequalis* individuals, nearby individuals originating from *Eriobotrya japonica* and could represent another *forma specialis*.

**Figure 3:**
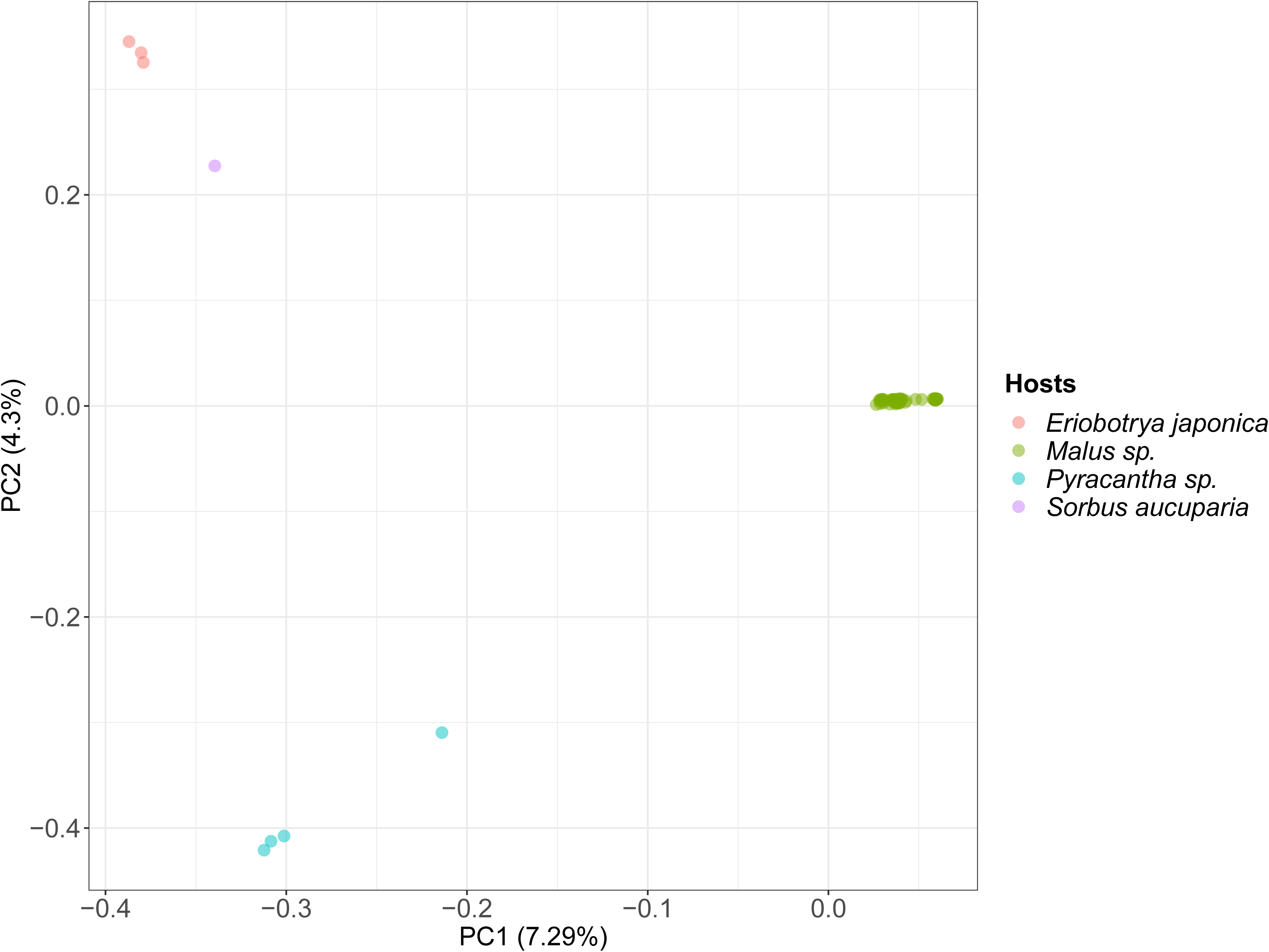
Population structure of *Venturia inaequalis*. PCA of the 71 *V. inaequalis* strains sampled on *Malus* spp. (green), *Eriobotrya japonica* (in red), *Pyracantha* sp. (blue) and *Sorbus aucuparia* (purple). The first axis differentiates strains sampled on apple trees from the others. Second axis separates strains sampled on Pyracantha sp. from those sampled on *E. japonica* and *S. aucuparia*.

### Gene predictions

Most of *V. inaequalis*, *V. pirina* and *V. aucupariae* strains contain around 11,600 predicted genes, while *V. asperata* strains contain fewer predicted gene models, around 10,400 (Figure 1D and Table S1). Virulence of plant pathogens relies on the secretion of small-secreted proteins (SSPs) that are often rich in cysteine residues (Lo Presti *et al*., 2015). The number of predicted SSPs of less than 300 amino acids ranges between 527 and 756 in *V. inaequalis* and *V. pirina* strains, representing between 4.8% and 6.1% of the predicted proteome (Figure 1D and Table S1). The PacBio assembly of the *V. inaequalis* EU-B04 isolate certainly explains the much higher number of predicted SSPs and percentage of the proteome. Despite a similar total number of predicted genes, the *V. aucupariae* isolate contains fewer SSPs (461; Figure 1D). In addition to a slightly reduced gene complement, *V. asperata* strains contain fewer predicted SSPs that represent less than 4% of the predicted proteome (between 395 and 427; Figure 1D and Table S1). Detailed analysis of the *V. inaequalis* and *V. pirina* predicted secretome identified putative race-cultivar and host-species specific SSPs (Deng *et al*., 2017). Similarly, in our report, differences in gene complement might explain the host range of the different species, especially the host specialization of *V. inaequalis formae specialis* and the low virulence of *V. asperata* on apple (Caffier *et al*., 2012). In the PacBio assembly, genes tend to be located within GC-equilibrated regions (Figure 2B and 2C). Although SSPs of *V. inaequalis* were reported closer to repetitive elements compared to other genes (Deng *et al*., 2017), SSPs do not tend to be located within long AT-rich regions as observed in *L. maculans* (Figure 2C and 2D; Rouxel *et al*., 2011).

### Invasion of transposable elements

The repetitive element content across all genomes was annotated using the REPET pipeline (Figure 1B; Table S1). The transposable element (TE) content covers 55.7% of the reference EU-B04 genome assembly, of which 66.4% are Class I LTR element, mainly assigned to Gypsy or Copia superfamilies (37.5% and 26.3%, respectively); and 9.5 % are Class II elements, mostly belonging to the TIR superfamily (8.2%) (Figure S1). Consistent with the exclusion of repeat content from CAM and CAP populations DNA libraries, only a median of 8.5% of the assemblies are covered with TEs (Figure 1B and Table S1). In other *V. inaequalis* genomes, the median coverage of TEs is 41.7% (Figure 1B and Table S1). TE coverage in the *V. aucupariae* assembly is similar, with 17.4% (Figure 1B and Table S1). In contrast, TE coverage is very limited in *V. pirina* (4.1% on average) and *V. asperata* (3.4% on average), consistent with exclusion of repeat content from DNA libraries of the former and limited number of TEs in the latter (Figures 1B and Table S1). The repetitive content of *V. asperata* strains differs from others as it does not contain Helitron and LINE elements, and is almost devoid of Copia elements (Figure S1 and Table S2). Gypsy and Copia TEs are predominant in all other *Venturia* strains. Notably Copia elements and class II TEs, especially TIR elements, appear to have greatly expanded after the divergence with *V. asperata* (Table S2 and Figure S1). Hierarchical clustering of the coverage of TE models annotated y REPET in all *Venturia* species suggests recent expansion of 32 Copia and Gypsy models (Table S3 and Figure S2); particularly two models (RLG-comp_VeinUTG-B-G309-Map3 and noCat_VeinUTG-B-R23-Map20) exhibits high coverage and are likely still actively transposing (Table S3 and Figure S2). A second cluster of 31 models might represent ancestral expansion of currently inactive TEs and a third cluster contains models with limited coverage (Figure S2). A few elements appear to be actively multiplying in certain genomes. For example, one Gypsy element (RLG-comp_VeinUTG-B-G1005-Map14) exhibits higher coverage in *V. asperata*, *V. aucupariae* and, to a lesser extent, in *V. pirina* compared to *V. inaequalis* strains (Table S3 and Figure S2), while another Gypsy element (RLG-comp_VeinUTG-B-R34-Map20) and a LARD element (RXX-LARD_VeinUTG-B-G646-Map11) show highest coverage in *V. inaequalis* 2199 and *V. inaequalis* B04 strains, respectively (Table S3 and Figure S2). In the *V. inaequalis* B04 reference genome, TEs are mostly located within AT-rich regions (Figure 2B and 2D), but many appear to be located in close vicinity to genes as reported for SSPs (Deng *et al*., 2017). In fungi, TE activity can be suppressed by Repeat-Induced Point mutation (RIP), a process that induces in repetitive DNA cytosine to thymine transition mutations during heterokaryon formation prior to meiosis (Watters *et al*., 1999). As such, RIP is a mechanism that produces AT-rich regions. Evaluation of RIP in *Venturia* strains showed that most TEs have been mutated through RIP (Figure 2E and Table S1). AT-rich regions in all strains are enriched in TpA di-nucleotides (Figure 2G), a feature typical of RIP in non-coding regions (Testa *et al*., 2016).

## Conclusions

Strains of the *Venturia* species complex are responsible for important diseases on Rosaceae fruit trees, as such they constitute an environmental and public health concern due to the massive use of fungicides necessary for their control. Our study provides valuable genomic resources for those interested in identification of molecular basis of *Venturia* species adaptation to their host and in mapping traits of interest (virulence, aggressiveness, resistance to fungicides etc…). In addition, these genomes will be used for comparative genomic studies to develop diagnostic molecular tools and to understand the evolution of genomes in relation to their hosts and in particular during the domestication of the apple tree.

## Supporting information

## Acknowledgments

The project was funded Plant Healt Department of INRA (ESCAPADES Project) and by the Agence Nationale pour la Recherche (Bioadapt ANR Gandalf ANR-12-ADAP-0009 and Fungisochore ANR-09-GENM-028 projects) and by the French Region Pays de la Loire (ROAD MOVIE project). JS and TP research was conducted in the framework of the regional program "Objectif Végétal, Research, Education and Innovation in Pays de la Loire“, supported by the French Region Pays de la Loire, Angers Loire Métropole and the European Regional Development Fund. LD was supported by the University of Angers, MDG was supported by Plant health Department of INRA and Region Pays de La Loire.

We thank Dr. P. Gladieux (INRA, Montpellier, France), Dr. XG Zhang (Shandong University, Taian, China), Dr. H Ishii (Kobe University, Kobe, Japan), Dr. Tim Roberts (East Malling Research, East Malling, England), Dr. A Kollar (JKI, Dossenheim, Germany) and Dr. E Dapena (SERIDA, Villaviciosia, Spain), Dr. F. Laurens (INRA, Angers, France) for collecting and providing strains or infected apple leaves.

This work was performed in collaboration with the GeT core facility (Toulouse, France; get.genotoul.fr), and was supported by France Génomique National infrastructure, funded as part of “Investissement d’avenir” program managed by Agence Nationale pour la Recherche (contract ANR-10-INBS-09).

The present work has benefited from the Cytometry facility of Imagerie - Gif, (www.i2bc.paris-saclay.fr), member of IBiSA (www.ibisa.net), supported by “France-BioImaging” (ANR-10-INBS-04-01), and the Labex “Saclay Plant Science”(ANR-11-IDEX-0003-02).

## Supplementary material

**Table S1**. Origin of sequenced strains and genome statistics

**Table S2**. Transposable element class coverage (bp as determined by REPET)

**Table S3**. Transposable element model coverage (bp as determined by REPET)

**Figure S1**. Genome coverage of classes of transposable elements (TEs) as determined by REPET in all *Venturia* strains, expressed as a percentage of the total TE content in each genome.

**Figure S2**. Hierarchical clustering of TE models annotated by REPET in all *Venturia* strains.

