## Supplementary material for "Population genome sequencing of the scab fungal species *Venturia inaequalis*, *Venturia pirina*, *Venturia aucupariae* and *Venturia asperata*"

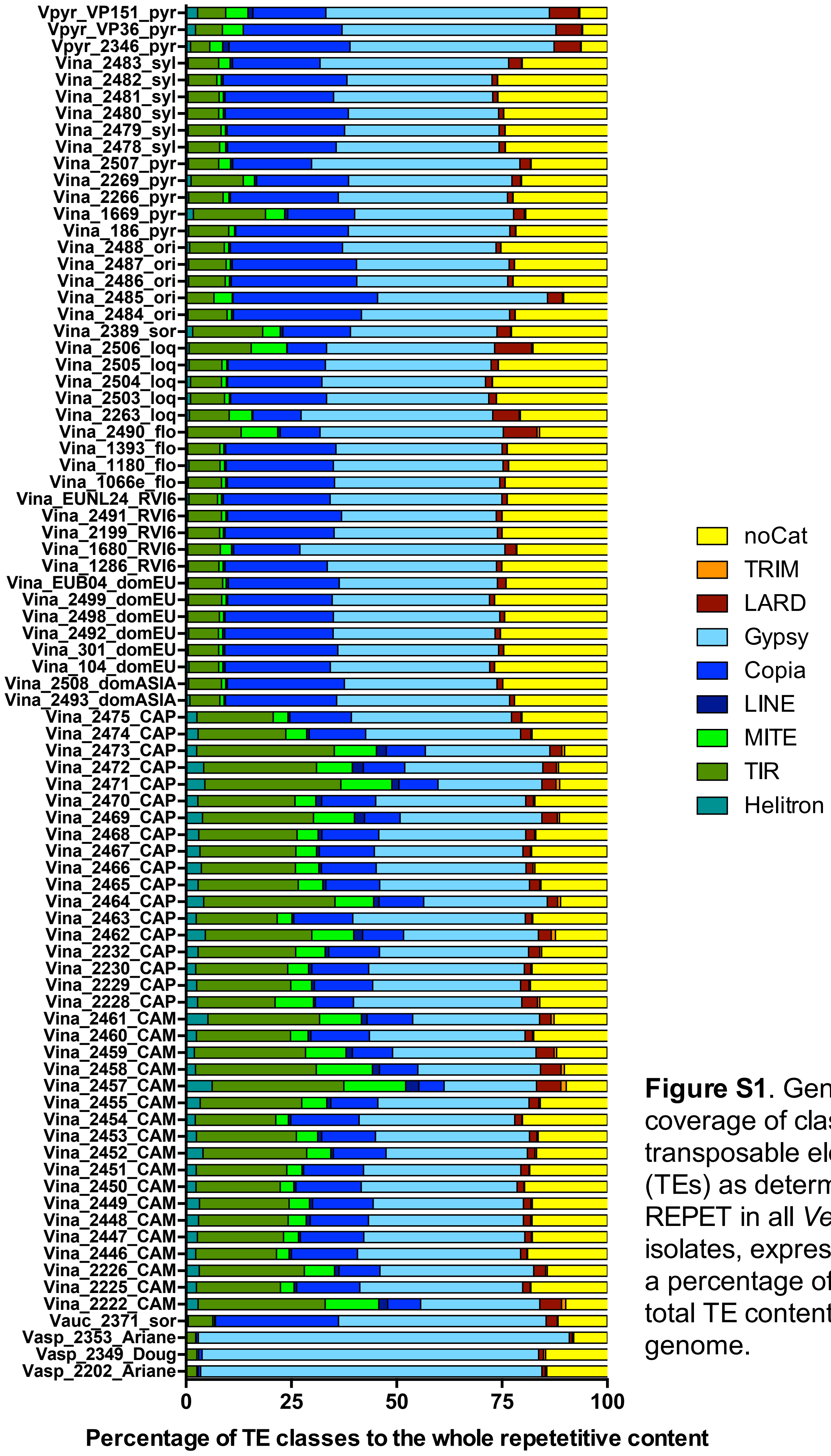


**Figure S1**. Genome coverage of classes of transposable elements (TEs) as determined by REPET in all *Venturia* isolates, expressed as a percentage of the total TE content in each genome.
