## Supplementary material for "Population genome sequencing of the scab fungal species *Venturia inaequalis*, *Venturia pirina*, *Venturia aucupariae* and *Venturia asperata*"

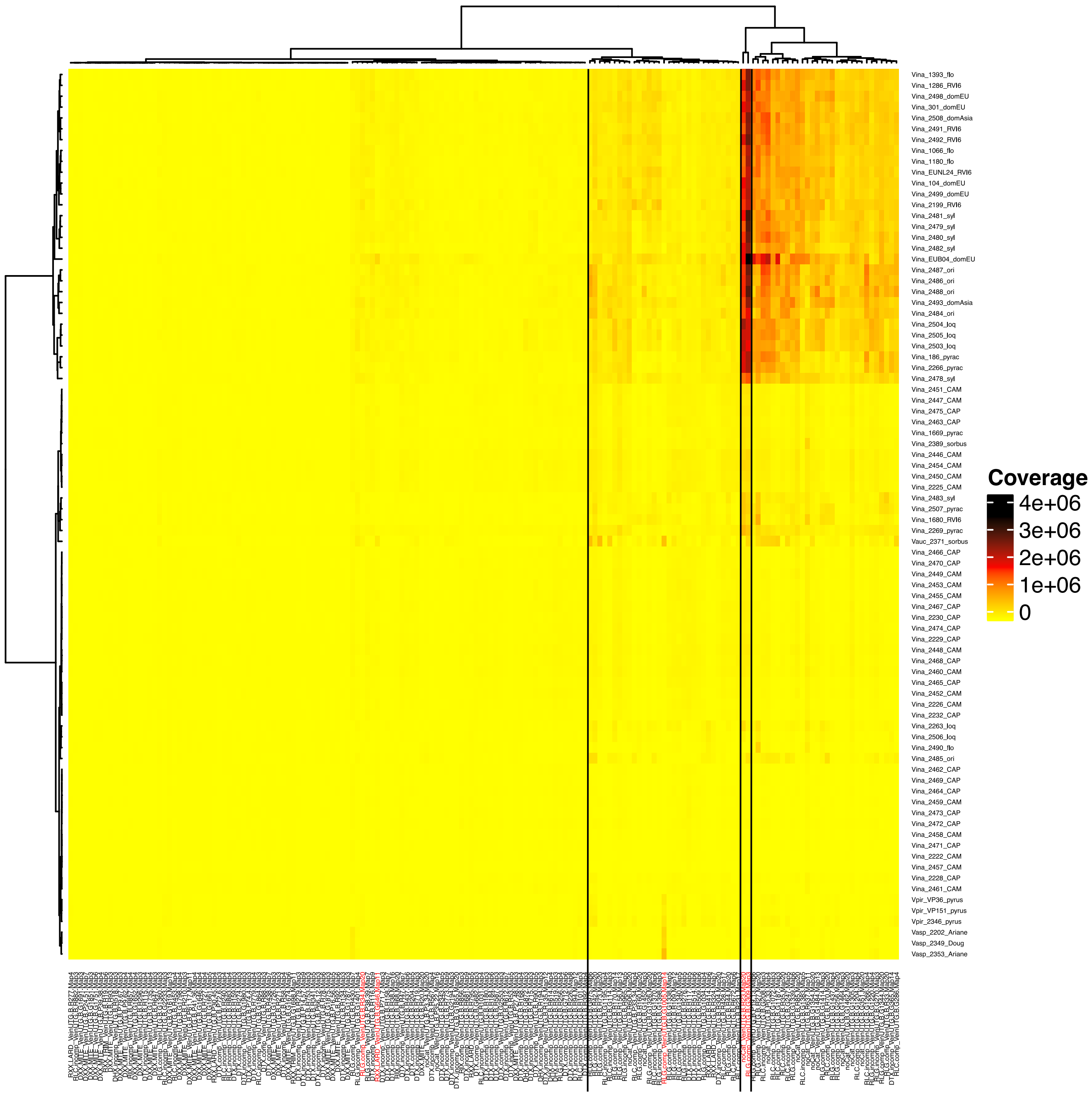


**Figure S2.** Hierarchical clustering of TE models annotated by REPET in all *Venturia* isolates. The Ward method with euclidian distance was employed. The color scale indicate the genome coverage (bp).
